# Biomechanical Implications of Spinopelvic Alignment on Femoral Head Cartilage and the Proximal Femoral Physis in Slipped Capital Femoral Epiphysis: A Theoretical Finite Element Analysis

**DOI:** 10.1101/2024.02.08.579521

**Authors:** Yogesh Kumaran, Muzammil Mumtaz, Carmen Quatman, Julie Balch-Samora, Sophia Soehnlen, Brett Hoffman, Sudharshan Tripathi, Norihiro Nishida, Vijay K. Goel

**Affiliations:** Engineering Center for Orthopedic Research (E-CORE), Departments of Bioengineering and Orthopaedic Surgery, University of Toledo, Toledo, OH, USA; Ohio State University Wexner Medical Center, Department and Orthopaedics, Columbus, OH, USA; Nationwide Children’s Hospital, Department of Orthopedics, Columbus, OH, USA; Yamaguchi University Hospital, Department of Orthopaedic Surgery, Ube, Yamaguchi, Japan

**Keywords:** Slipped capital femoral epiphysis, spinopelvic parameters, finite element analysis

## Abstract

**Background:** Slipped capital femoral epiphysis (SCFE) is a prevalent pediatric hip disorder. Recent studies suggest the spine’s sagittal profile may influence the proximal femoral growth plate’s slippage, an aspect not extensively explored. This study utilizes finite element analysis to investigate how different spinopelvic alignments affect shear stress and potential slippage at the growth plate.

**Methods:** A finite element model was developed from CT scans of a healthy adult male lumbar spine, pelvis, and femurs. The model was subjected to various sagittal alignments through rotational boundary conditions. Simulations of two-leg stance, one-leg stance, walking heel strike, ascending stairs heel strike, and descending stairs heel strike were conducted. Parameters measured included hip joint contact area, stress, and maximum Tresca (shear) stress on the growth plate.

**Findings:** Posterior pelvic tilt cases indicated larger shear stresses compared to the anterior pelvic tilt variants except in two leg stance. Two leg stance resulted in decreases in the posterior tilted pelvi variants compared to anterior tilted pelvi, however a combination of posterior pelvic tilt and high pelvic incidence indicated larger shear stresses on the growth plate. One leg stance and heal strike resulted in higher shear stress on the growth plate in posterior pelvic tilt variants compared to anterior pelvic tilt, with a combination of posterior pelvic tilt and high pelvic incidence resulting in the largest shear stress.

**Interpretation:** Our findings suggest that posterior pelvic tilt and high pelvic incidence can lead to increased shear stress at the growth plate. Activities performed in patients with these alignments may predispose to biomechanical loading that shears the growth plate, potentially causing slippage.

## Introduction

Slipped capital femoral epiphysis (SCFE) is a multifactorial pathology of the hip commonly affecting children between the ages of 8 and 15 years with an incidence of 0.3-2/100,000 [1]. Early identification of high-risk patients for SCFE is crucial for implementing preventive strategies against slippage at the proximal femoral physis and its adverse sequelae [2]. Obesity, endocrinopathies, and vitamin D deficiency are risk factors that have been implicated in its development [3, 4]. Biomechanical factors, including acetabular version, repetitive weight-bearing activities, sports, and other physical exertions, also contribute to increased shear stresses at the proximal femoral physis, leading to slippage [5, 6]. Recent studies have also pointed to the sagittal profile of the spinopelvic complex as a contributing factor to growth plate shear and slippage [7, 8].

Sagittal balance refers to the physiologic alignment of the spine which maximizes kinetic efficiency and minimizes mechanical stress. Balance exists when an individual’s weight is positioned on a vertical axis aligned slightly posterior to the rotational axis of the femoral heads [9, 10]. Spinopelvic parameters determine the overall sagittal balance of the spinopelvic complex and include lumbar lordosis (LL), sacral slope (SS), pelvic tilt (PT), and pelvic incidence (PI) (Figure 1) [11–14].

**Figure 1:**
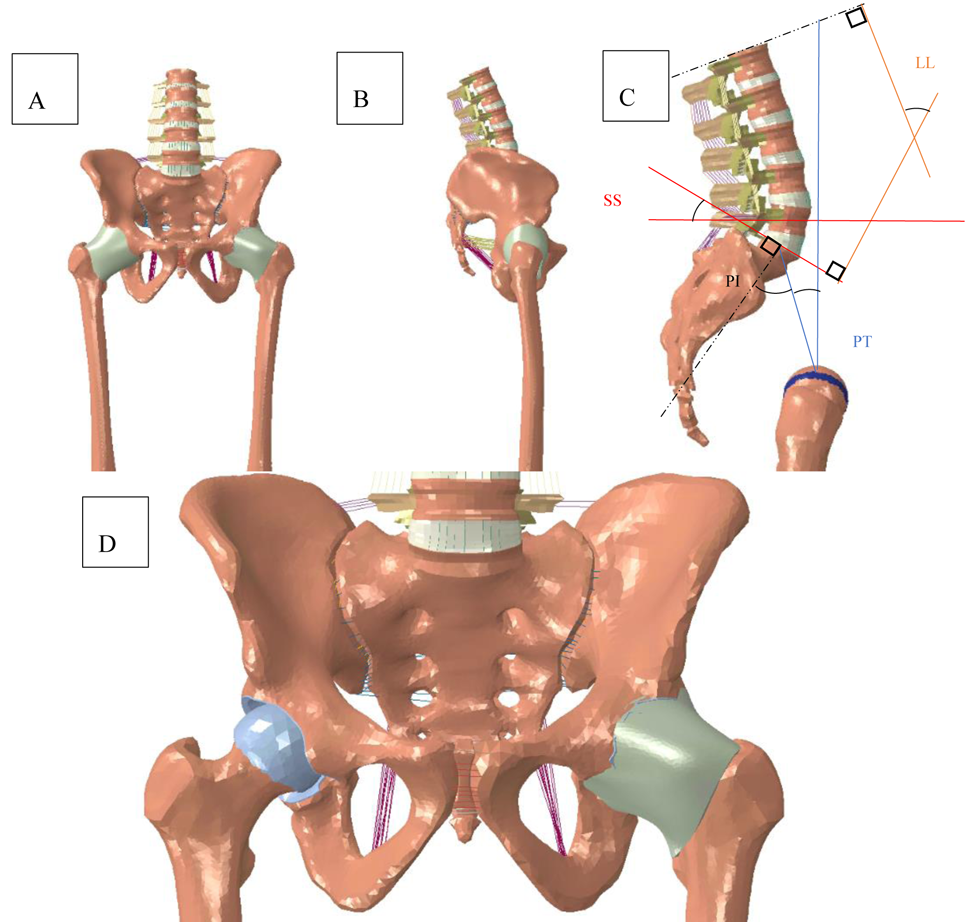
Finite element model with the following spinopelvic parameters: PI = 46, SS = 26.7, PT = 20, LL = 51. A) Represents the anterior view of the model, B) represents the sagittal profile, C) represents the spinopelvic angles (PT, PI, SS, LL), D) represents the articulating surface of the hip joint with the capsule removed (right hip) and the capsule surrounding the hip joint (left hip).

While the relationship between hip and spine mechanics has been extensively studied in adults, its impact on the SCFE population remains less explored [15, 16]. Controversy exists on whether PI and other spinopelvic parameters contribute to SCFE. A cadaveric study performed by Gebhart et al identified smaller PI angles in post-SCFE deformity patients compared to normal specimens [7]. On the other hand, a retrospective study performed by Wako et al determined no relationship between PI and SCFE [8]. These discrepancies highlight the need for biomechanical analyses to understand the role of load distribution on the femoro-acetabular joint and proximal femoral physis due to variations in sagittal spinopelvic parameters.

To the best of the authors’ knowledge, this is the first biomechanical study to examine SCFE encompassing the complete spinopelvic complex, including the femur, hip joint, pelvis, and spine [17–21]. We hypothesize that sagittal spinopelvic alignment influences stress redistribution on the femoral head and proximal femoral physis. This study utilizes a theoretical finite element (FE) model to investigate the contribution of sagittal spinopelvic parameters to SCFE biomechanics.

## Methods

### Original FE model creation

A non-linear, ligamentous FE model was developed using computed tomographic (CT) scans of a healthy adult lumbar spine, pelvis, and femur with no abnormalities, deformities, or severe degeneration. The CT scans were reconstructed and segmented using MIMICS software (Materialize Inc., Leuven, Belgium). Mesh was applied to the reconstructed geometry using Meshlab Open-Source Software [22].

Abaqus 2019 (Dassault Systèmes, Simulia Inc., Providence, RI, USA) was used to assemble the meshed components and perform the subsequent analyses. All geometries were imported and the “tri to tet” feature was used to convert the triangular shell surface mesh into solid tetrahedral mesh (C3D4). The vertebral bodies, pelvis and femurs were modelled with a 0.5-1mm cortical bone shell containing a core of cancellous bone. The annulus fibrosa was simulated as a composite solid containing alternating ±30° collagen fibers modelled as rebar elements using the “no compression” property. The nucleus pulposa was simulated as a linear elastic material. The facet joints were modeled using three-dimensional gap elements with a defined clearance of 0.5mm. The ligamentous structures of the spine and sacroiliac joints were modelled as truss elements. Table 1 lists the material properties used in the FE models [23–26]. Validation of the vertebral components of the lumbar spine are described elsewhere [25–30].

**Table 1:**
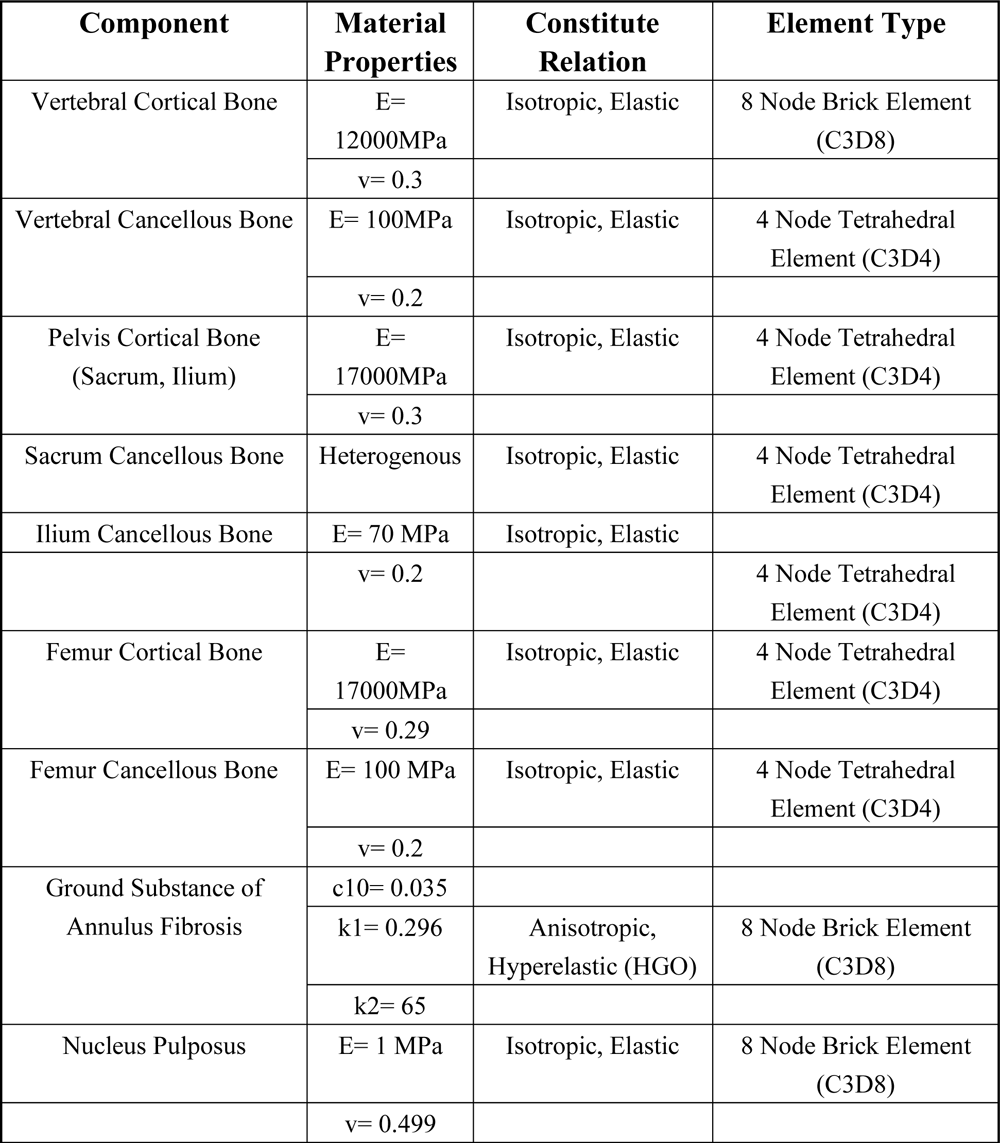
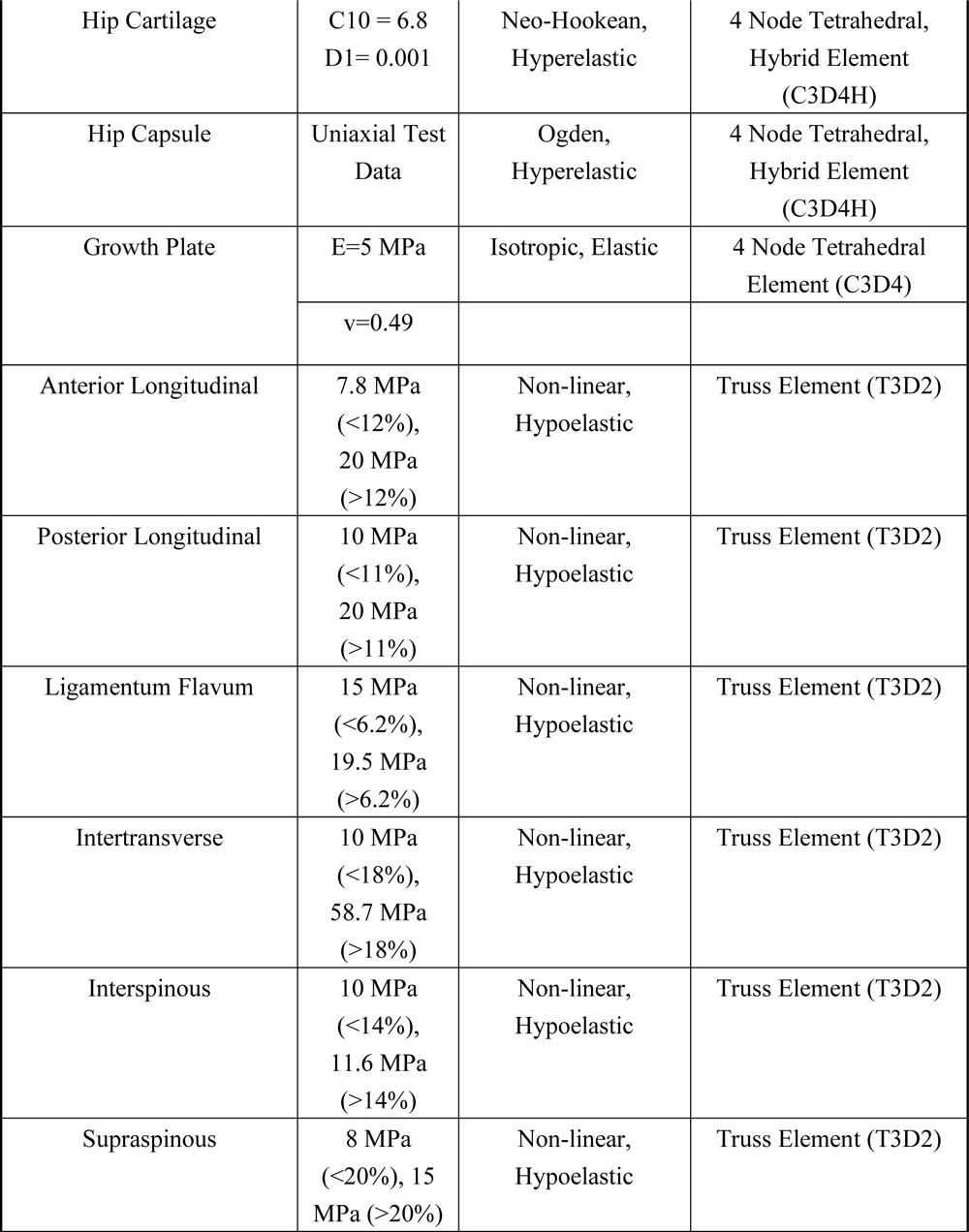
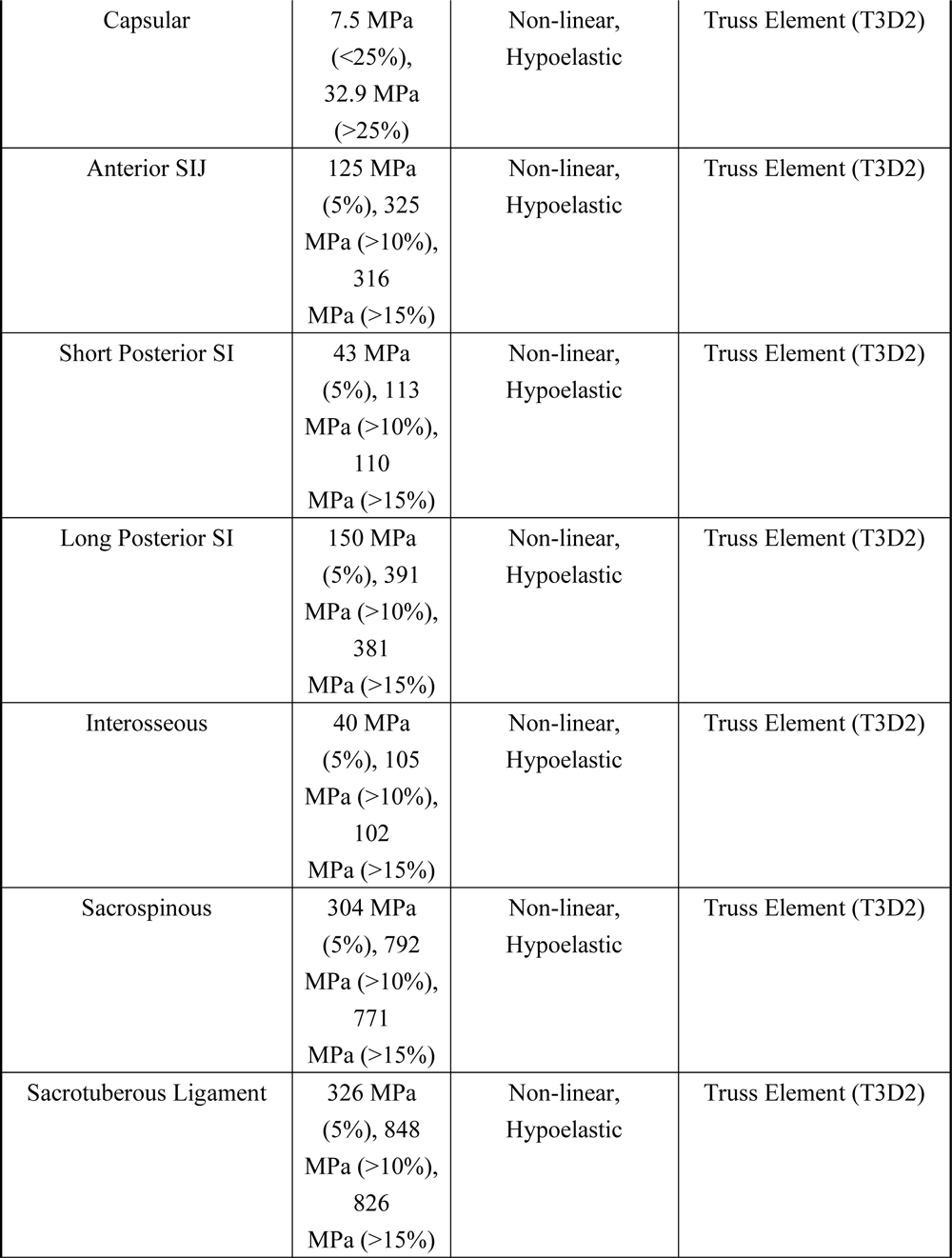
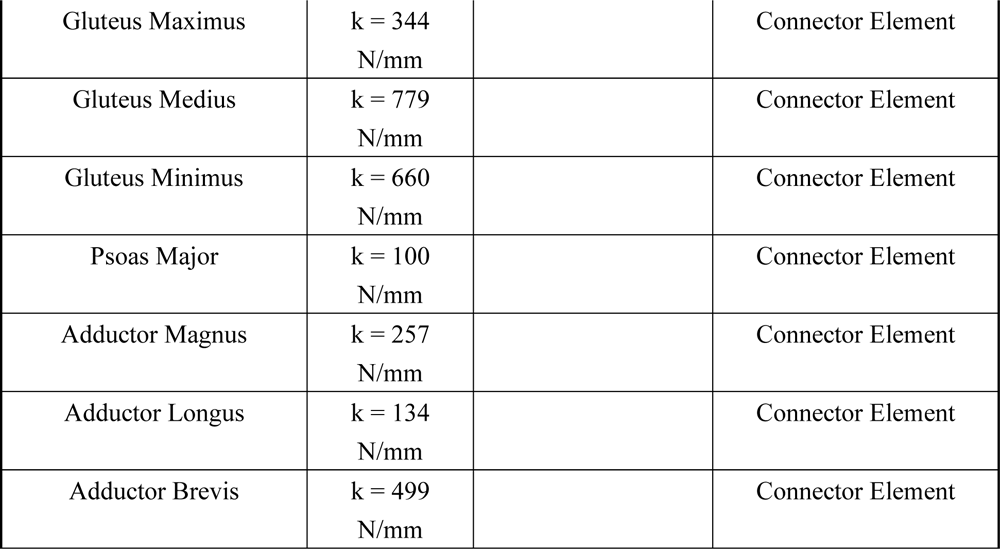
Material properties assigned to the finite element models [17, 21, 27, 34, 36, 37, 39, 51, 62–65]. E represents Young’s modulus, v represents Poisson’s ratio.

### Hip joint creation and validation

The femoral heads were positioned relative to the pelvis according to anatomical parameters described by Wu et al [31]. Femoral head and acetabular cartilage were modelled by adding a 1mm layer of mesh with hybrid formulation (C3D4H) applied on both surfaces [32]. Ten to twelve random nodal coordinates of the femoral heads and acetebulae were obtained to create a best fit sphere whose centroid coordinates were used to properly place the femoral heads within the acetabulae [33]. The cartilage was modelled with incompressible, neo-Hookean hyperelastic material properties [34]. The hip joint was adjusted until the joint space fell within reported parameters determined by Goker et al (3.43± 0.40 for the right hip and 3.48±0.68 for the left hip) [35]. The hip joint capsule was created in Solidworks 2022 (Dassault Systèmes, Simulia Inc., Providence, RI, USA) based on circumferential locations defined by Stewart et al [36]. Hyperelastic material properties were applied to this geometry based on the Ogden strain energy potential and utilizing uniaxial test data based on a study performed by Stewart et al [36]. Muscles spanning the hip joint from their physiologic origins and insertions were included in the FE model as connector elements with stiffnesses based on Phillips et al (Table 1) [37].

To assess and validate proper functionality of the hip joints, contact stresses and areas were evaluated based on simulated positions corresponding to two-leg stance (2LS), walking heel strike (WHS), ascending stairs heel strike (AHS), and descending stairs heel strike (DHS) per femur angles based on Bergmann et al’s data [38, 39]. Peak and average contact stresses (MPa) and contact area (mm^2^) at the hip joint were validated against previous studies [34, 39, 40].

The growth plate was modelled by sectioning 7mm of the femoral head [41]. The right and left femur physeal-diaphysis angle (PDA) was 39° and 44°, respectively (Figure 2) [20]. Elastic material properties were applied to the growth plate (Table 1) [17, 20, 21, 42]. The original model had the following spinopelvic parameters: SS=31.7°, PT=9.8°, PI=41.5°, and LL=46°.

**Figure 2:**
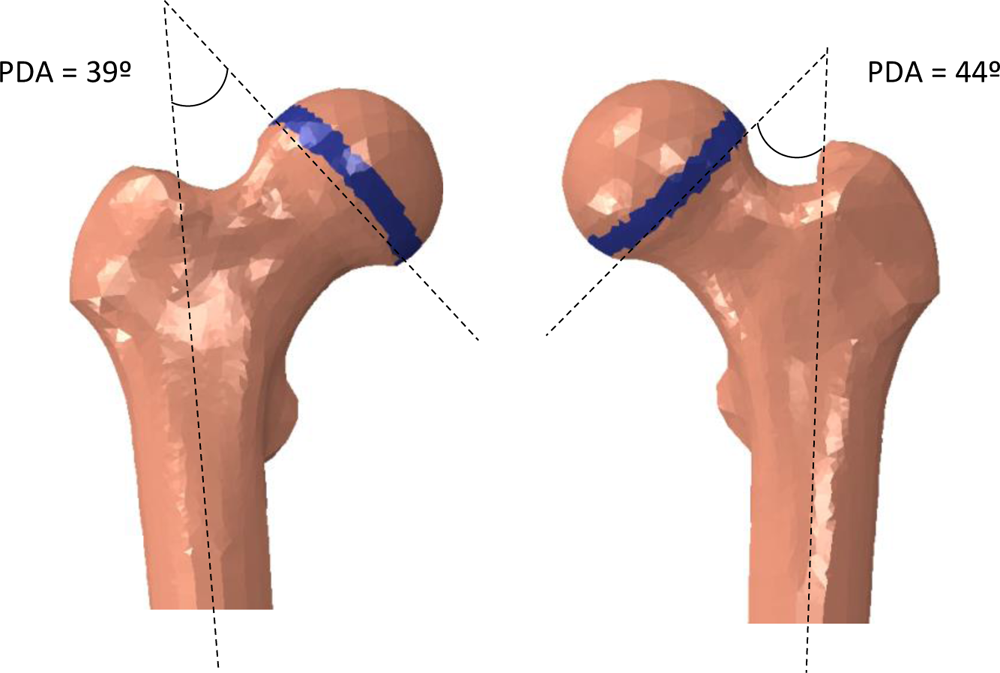
Representation of the right and left femurs with 7mm thick growth plates. The PDA for the right and left femurs were 39° and 44°, respectively. The PDA was measured by drawing a line through the intremedullary canal in the femur and an intersecting line parallel to the base of the epiphyseal growth plate.

This model was modified with various SS and PT angles by rotating the sacrum and pelvis, respectively [43]. The SS angles were incrementally increased by 5° to obtain PIs of 36°, 41°, 46°, and 51°, respectively. To understand the influence of PT on the growth plate, each SS modified model had two variants: high PT (posteriorly tilted) and low PT (anteriorly tilted) cases (Table 2).

**Table 2:** Spinopelvic parameters of the finite element models. Low PT refers to anteriorly tilted pelvi, whereas high PT refers to posterior pelvic tilt.

| **Model** | **Sacral Slope**<br>**(SS°)** | **Pelvic Tilt**<br>**(PT°)** | **Lumbar**<br>**Lordosis**<br>**(LL°)** |
| --- | --- | --- | --- |
| PI 36° Low PT | 26.7 | 9.5 | 41 |
| PI 36° High PT | 16.7 | 19.5 | 41 |
| PI 41° Low PT | 31.7 | 9.8 | 46 |
| PI 41° High PT | 21.7 | 19.8 | 46 |
| PI 46° Low PT | 36.7 | 10 | 51 |
| PI 46° High PT | 26.7 | 20 | 51 |
| PI 52° Low PT | 41.7 | 10.4 | 56 |
| PI 52° High PT | 31.7 | 20.4 | 56 |

### Loads, boundary conditions, and analysis

A 400 N compressive follower load was applied following the curvature of the L1-L5 vertebrae through wire elements to simulate the passive effect of muscle forces and weight of the upper trunk [39]. A 500 N force was distributed between the sacral promontory and pubic symphisys to simulate body weight on the pelvis and femur [39]. Two leg stance (2LS), right leg one leg stance (1LS), AHS, DHS, and WHS cases were simulated to evaluate various activities’ impact on the femoral heads and growth plate. Boundary conditions were applied at the base of the femurs to fix the model. Average contact stress and contact area of the femoral head was calculated for each motion and alignment. To evaluate shear stresses on the proximal femoral growth plate, maximum Tresca stresses were recorded.

## Results

Validation was performed on the model to confirm functionality of the hip joints. Average contact stresses were found to be in range of previous studies with peak contact stresses being slightly higher (Tables 3, 5 and 6). Regarding contact areas, values were in-line of previous studies (Tables 4 and 7).

**Table 3:** Hip validation for peak and average contact stresses [39].

| Motion | Joukar et al |  |  |  | Current FE Model |  |  |  |
| --- | --- | --- | --- | --- | --- | --- | --- | --- |
|  | Average Contact stress (Mpa) |  | Peak Contact stress (Mpa) |  | Average Contact stress (Mpa) |  | Peak Contact stress (Mpa) |  |
|  | Right | Left | Right | Left | Right | Left | Right | Left |
| Standing | 1.81 | 1.87 | 6.2 | 5.96 | 1.77 | 1.875 | 6.253 | 5.983 |
| Flexion | 1.65 | 2.29 | 5.28 | 5.48 | 1.97 | 2.15 | 6.829 | 6.946 |
| Extension | 1.86 | 1.76 | 7.4 | 5.3 | 1.78 | 1.77 | 5.686 | 5.268 |
| RB | 1.94 | 1.62 | 7.1 | 4.8 | 1.91 | 1.81 | 6.783 | 5.582 |
| RR | 1.66 | 1.94 | 6.1 | 6 | 1.79 | 1.93 | 6.126 | 6.131 |

**Table 4:** Hip validation results for contact area [39].

| Motion | Joukar et al |  | Current FE Model |  |
| --- | --- | --- | --- | --- |
|  | Average Contact area (mm2) |  | Average Contact area (mm2) |  |
|  | Right | Left | Right | Left |
| Standing | 224 | 207 | 218.977 | 224.105 |
| Flexion | 232 | 222 | 216.146 | 258.874 |
| Extension | 239 | 213 | 208.619 | 207.565 |
| RB | 238 | 201 | 214.363 | 217.741 |
| RR | 224 | 207 | 217.139 | 217.586 |

**Table 5:**
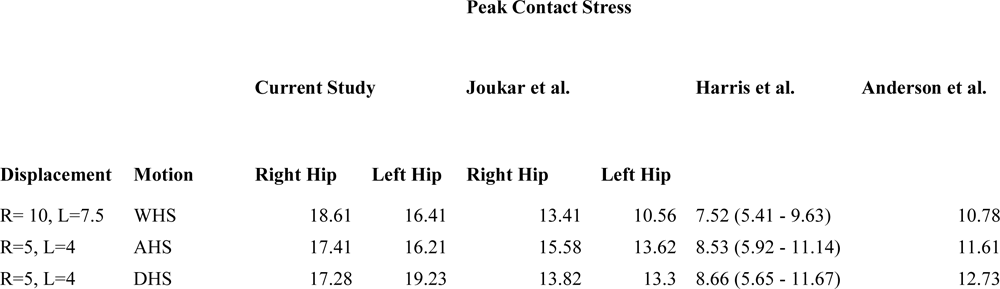
Hip validation for peak contact stresses in various heal-strike scenarios [34, 39, 40].

| Peak Contact Stress |  |  |  |  |  |  |  |
| --- | --- | --- | --- | --- | --- | --- | --- |
|  |  | Current Study |  | Joukar et al. |  | Harris et al. | Anderson et al. |
| Displacement | Motion | Right Hip | Left Hip | Right Hip | Left Hip |  |  |
| R= 10, L=7.5 | WHS | 18.61 | 16.41 | 13.41 | 10.56 | 7.52 (5.41 - 9.63) | 10.78 |
| R=5, L=4 | AHS | 17.41 | 16.21 | 15.58 | 13.62 | 8.53 (5.92 - 11.14) | 11.61 |
| R=5, L=4 | DHS | 17.28 | 19.23 | 13.82 | 13.3 | 8.66 (5.65 - 11.67) | 12.73 |

**Table 6:**
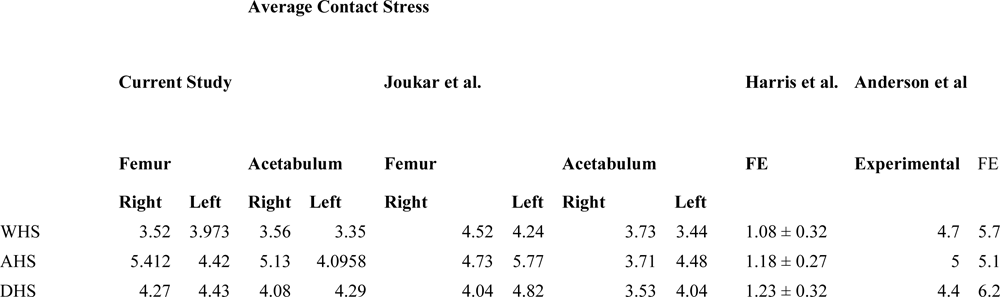
Hip validation results for average contact stresses in various heal strike scenarios [34, 39, 40].

**Table 7:**
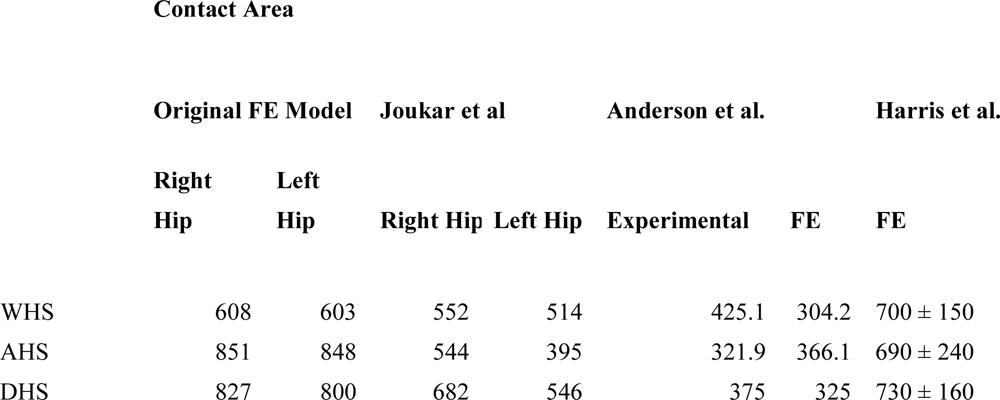
Hip validation results for average hip contact area in various heal strike scenarios [34, 39, 40].

Overall, higher PT indicated larger contact areas compared to low PT variants. A combination of higher PT and PI resulted in substantially larger contact areas compared to low PT and low PI for each simulated motion (∼18% increases) (Figure 3).

**Figure 3:**
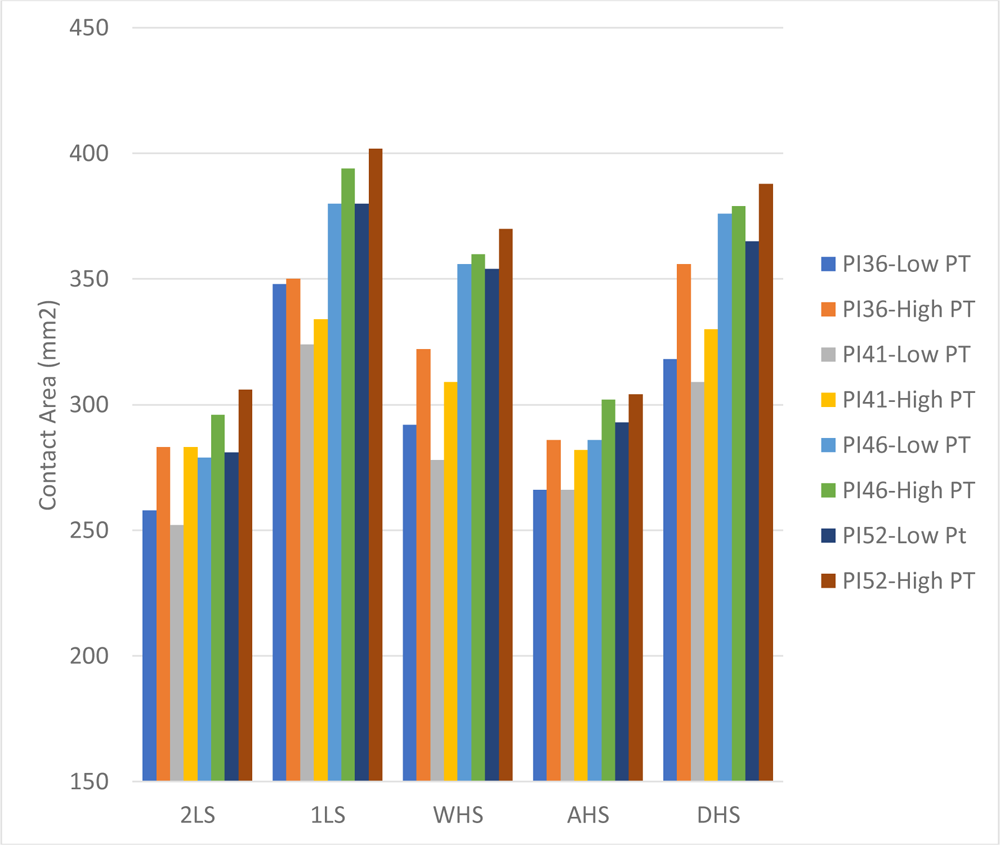
Contact area (mm^2^) on the femoral head cartilage in various stances. 2LS refers to two leg stance, 1LS refers to one leg stance, WHS refers to walking heel-strike, AHS refers to ascending stairs heel strike, and DHS refers to descending stairs heel strike.

Regarding hip contact stresses, the heel strike simulations (WHS, AHS, and DHS) resulted in the largest values compared to 2LS and 1LS, especially in the PI41 cases. The 1LS and 2LS simulations indicated that as PT increased, larger contact stresses were mitigated (Figure 4). All heel strike simulations showed a similar trend with higher contact stresses in the high PT variants (∼10% increases), except for WHS. In WHS, PI41-High PT resulted in the largest contact stress with marginal changes in higher PIs. AHS indicated lower contact stresses in PI52 compared to all other cases in AHS, with PI46 indicating the largest contact stress. In DHS, PI41 indicated the largest contact stresses compared to the other cases.

**Figure 4:**
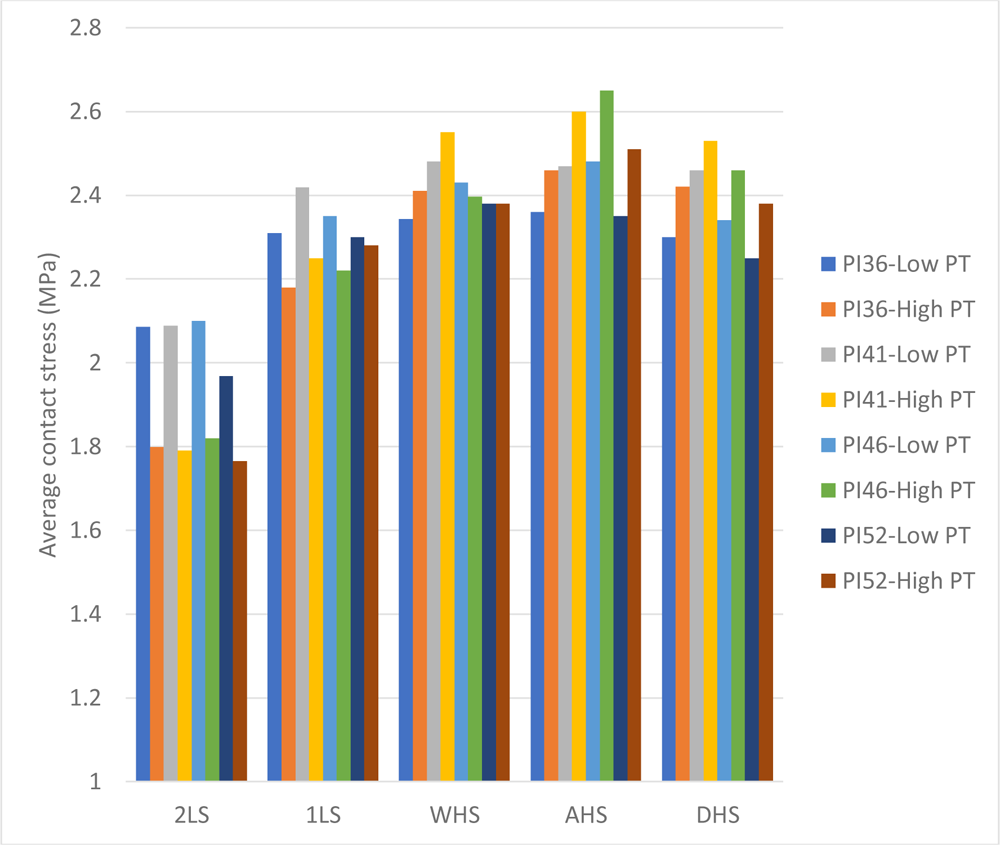
Average hip contact stress (MPa) on the femoral head cartilage in various stances.

Regarding Tresca stresses, heel strike simulations resulted in the largest stresses on the growth plate with AHS indicating the largest values overall. High PT variants resulted in larger shear stresses compared to the low PT variants (∼18% increases) except in 2LS. A combination of high PT and PI resulted in the largest overall Tresca stress on the growth plate for each simulated case (10-24% increases) (Figure 5). In 1LS, WHS, AHS and DHS, Tresca stresses for the high PT variant of PI36 was comparable to PI46-High PT, though smaller than PI52-High PT.

**Figure 5:**
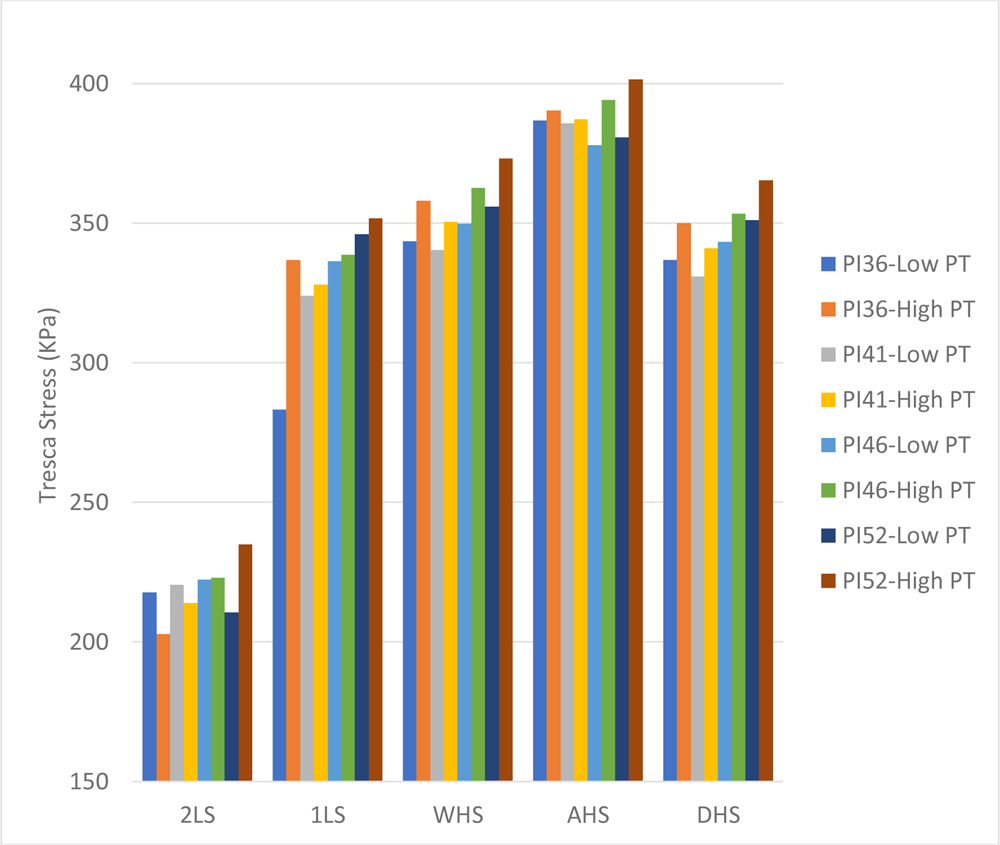
Maximum Tresca (shear) stress in KPa on the growth plate in various stances.

## Discussion

The aim of the current study was to elucidate the role of spinopelvic parameters on the hip joint and an open femoral growth plate to assess the potential for development of SCFE. Current research in this area is limited, with only a few studies leveraging FE models to examine SCFE under different loading scenarios [17–21, 42]. Our findings revealed that a combination of posterior pelvic tilt (PT) and high pelvic incidence (PI) angles lead to large hip contact area and elevated shear stresses on the proximal femoral physis.

Previous reports have stated that higher pelvic incidence and sacral slope angles contribute to larger mechanical stresses on the hip joint [43, 44]. Lazennec and colleagues described clinical consequences of sagittal imbalance in pelvic and sub-pelvic regions in patients and eluded to how atypical postures and morphology contribute to disturbances in the hip joint [15, 16]. Disruption of the spinopelvic complex displaces mechanical forces and increases load absorption on the intervertebral discs and femoroacetabular joints. The present study suggests that various spinopelvic alignments distribute stresses at the hip joint and growth plate differently, specifically patients with high pelvic incidence and posterior pelvic tilt potentially being more prone to slippage in SCFE [6]. Conflicting results are present in the current literature regarding PI and SCFE. An osteological, adult cadaveric study performed by Gebhart et al identified lower PI in post-SCFE specimens compared to a control of normal specimens [7] [45, 46]. Our results indicate that lower pelvic incidence angles contribute to larger contact stress at the hip joint, and seldom to shear stresses on the epiphyseal growth plate, which are two components that are risk factors for SCFE [6]. Additionally, the type of activity performed and degree of pelvic tilt also determine the overall shear at the growth plate. For instance, some postures indicated that low pelvic incidence combined with posterior pelvic tilt (PI36-HighPT) contributed to shear at the growth plate more than higher pelvic incidence (PI46). However, in most postures, high pelvic incidence alone and posterior pelvic tilt combined with high pelvic incidence resulted in the largest shear at the growth plate. This finding can be attributed to the high sacral slope and lumbar lordosis angles distributing a shear load through the femur and proximal femoral growth plate. Contrast this to the low pelvic incidence cases where the model remains more vertical due to a low sacral slope, the load distributing from the lumbar spine through the femoro-acetabular joint and growth plate were likely compressive forces rather than shear.

A retrospective observational study by Wako et al did not identify a relationship between PI and SCFE, though determined a significance in retroverted acetabuli and excessive coverage of the anterior and superior acetabulum in SCFE patients compared to a control group [8]. Current studies have mixed results on whether SCFE-affected hips are associated with the phenomena of anteversion or retroversion [7, 8, 47, 48]. Additionally, many studies do not correct for pelvic tilt which has the potential to overestimate acetabular version [49, 50]. In the current study, two pelvic tilt variants were examined for posterior and anterior pelvic tilt. Results indicated high contact area in the posterior pelvic tilt variants compared to anterior tilt which can be attributed to larger coverage of the posterior femoral head with the superior acetabulum. Contact stresses resulted in the same trend in 2LS and 1LS, though the opposite was seen in walking, ascending stairs, and descending stairs as posterior tilted variants indicated higher contact stress. Since stress is defined as the force applied over a certain area, this may seen counterintuitive, though as described by Henak et al, contact location, distribution and direction change during walking, ascending stairs, and descending stairs. In our case, the anterior tilted pelvi along with the angles of the femur in the heel strike simulations distributed the load anteriorly and superiorly compared to posterior tilt, thus agreeing with Henak and colleagues’ findings [51]. Our results suggest that patients may experience higher contact stress on the femoral head cartilage during gait, especially in patients with excess posterior pelvic tilt [8, 49, 51–53]. Along with higher femoral head contact stress, patients with posterior pelvic tilt may be more prone to slip due to higher shear stresses at the growth plate.

In the adult population, atypical pelvic postures produce consequences to the stability of the hip after arthroplasty, contributing further to imbalance of sagittal spinopelvic parameters [15, 16]. Furthermore, high pelvic incidence has been suggested to contribute to excessive loading on the femoral head cartilage leading to osteoarthritis of the hip [44, 54, 55]. Regarding SCFE, osteoarthritis has remained a problematic sequealae in young adult patients who experienced severe slip or surgical pinning to repair the proximal femoral physis [56, 57]. Long term follow-up of these patients commonly exhibit hip cartilage degeneration due to CAM deformity and femoroacetbular impingement [58, 59]. Based on these previous findings and the current study’s results, a pediatric patients dynamic mechanical history, surgical history of pinning, and sagittal alignment may suggest a compounding deleterious effect on the hip cartilage and early-onset osteoarthritis in sagittally imbalanced patients [60]. Future clinical and biomechanical studies will further explore this topic.

The current study’s finite element (FE) model, while insightful, is subject to certain limitations. Notably, we employed an adult model in this theoretical analysis. This decision stemmed from the fact that CT imaging for SCFE patients often focuses solely on the pelvic area, excluding critical details like lumbar lordosis and sacral slope. Conducting additional spine-focused CT scans would unnecessarily expose children to extra radiation. Consequently, crafting patient-specific models encompassing the complete spinopelvic complex with varied spinal alignments in pediatric SCFE patients necessitated certain simplifications. While it’s recognized that pelvic and spinal morphology evolve from childhood through adulthood, the study’s results are deemed reliable and indicate a potential mechanical linkage between spinopelvic parameters and the pediatric growth plate. The absence of validated pediatric spine models in the literature, predominantly due to the lack of pediatric cadaver studies, and the prevalent reliance on scaling methods for spine modeling in children, further justified our approach Furthermore, while the reference range for pelvic incidence (PI) in adults spans from 33 to 85, our study faced a technical limitation in that our finite element (FE) model could only simulate PIs between 36 and 51 [61]. This restriction was due to element penetration and contact errors encountered when attempting to adjust the PI beyond 51. As a result, the range of PIs explored in our study was narrower than ideal. This constraint potentially limits the generalizability of our findings across the full spectrum of PI values typically observed in clinical settings. Additionally, muscle forces spanning the hip and spine were simplified as passive connector elements with elastic properties and follower loads. Global mechanics were of interest to the authors rather than micro-mechanics, therefore the growth plate was simulated as a linear, elastic material property [17, 21, 42]. Additionally, the physeal-disphyseal angle and physeal thickness were held constant due to variation of plate thickness and angle having little contribution to stresses on the growth plate [17, 18, 20]. Lastly, only a male model was constructed in this study. The biomechanics of female pelvi may distribute loads differently compared to male pelvi, therefore future studies should examine sexual dimorphism in pediatric pelvi.

This study suggests that patients with specific sagittal alignments such as low pelvic incidences in some activities and high pelvic incidences combined with posterior tilted pelvi may be more prone to slip due to changes in contact stresses at the femoral head and high shear stress on the proximal femoral growth plate. It may be necessary for hip preservation surgeons to consider sagittal imbalance as a potential risk factor for SCFE, though future studies are required to confirm this theory.

## Supplementary Data

